# Artificial Intelligence driven Benchmarking Tool for Emission Reduction in Canadian Dairy Farms

**DOI:** 10.1101/2024.05.29.596461

**Authors:** Pratik Mukund Parmar, Hangqing Bi, Suresh Neethirajan

**Affiliations:** Faculty of Computer Science, 6050 University Avenue, Dalhousie University, Halifax, Canada; Faculty of Agriculture, Agricultural Campus, PO Box 550, Dalhousie University, Truro, NS, B2N 5E3

**Keywords:** Digital agriculture, Smart farming, climate change, dairy emissions, Canadian dairy, dairy cows, livestock production, Precision Livestock Farming, Data Analytics

## Abstract

This study develops an Artificial Intelligence-driven benchmarking tool to reduce methane emissions in Canadian dairy farms, responding to the urgent need to mitigate environmental impacts from agriculture. Utilizing a comprehensive dataset from over 1000 dairy farms and processors across Canada, combined with satellite-driven methane emission data, we apply advanced machine learning technologies and data analytics, including geospatial analysis and time series forecasting. This approach identifies critical emission hotspots and temporal trends. We tested several predictive models—ARIMA, LSTM, GBR, and PROPHET—with the LSTM model showing the greatest accuracy in forecasting emissions, demonstrated by the lowest Root Mean Squared Error (RMSE) of 15.40. Our results highlight the transformative potential of AI tools in agricultural environmental management by providing dairy farmers and policymakers with precise, real-time emission insights. This facilitates informed decision-making and the implementation of effective emission reduction strategies. This study not only advances understanding of emission dynamics in dairy farming but also underscores the role of technology in sustainable agricultural practices and achieving environmental targets consistent with global agreements.

## Introduction

The dairy industry is an integral component of Canada’s agricultural sector, significantly bolstering both the economy and food security. With over 9,400 dairy farms and numerous processing facilities, this sector is pivotal in the agricultural framework (AAFC, 2023). However, it also poses environmental challenges, primarily due to its considerable emissions of methane, a potent greenhouse gas with a global warming potential 28 times that of carbon dioxide over a century (Yoro and Daramola, 2020).

The primary source of methane emissions in dairy farming is enteric fermentation, a natural digestive process in ruminants such as cows and buffaloes. This process is compounded by the decomposition of manure in storage facilities, which significantly amplifies these emissions, thereby contributing to climate change, air pollution, and ecological degradation (Skytt et al., 2020). Addressing these emissions is critical for Canada to meet its international environmental commitments, including those under the Paris Agreement.

Traditional methods of monitoring and mitigating these emissions have predominantly relied on ground-based measurements, including assessments of dairy cow feeding, ruminant behavior, and manual on-farm animal husbandry data collection (Laubach et al., 2024). The monitoring and mitigation strategies in the dairy industry have traditionally involved direct measurements and manual data collection, which were not only time-consuming. The on-farm methane emissions measurement approaches were labor-intensive, error-prone, and lacked the necessary resolution to effectively capture the complex dynamics of emissions across diverse farming operations.

The advent of satellite-based remote sensing technology, coupled with advanced data analytics, has transformed the ability to monitor emissions (Zhang et al., 2023). Recent advances in technology have introduced satellite-based remote sensing as a valuable tool for monitoring greenhouse gas (GHG) emissions from various sources, including dairy farms (Palmer et al., 2021). Satellites such as the Greenhouse Gases Observing Satellite (GOSAT) and the Tropospheric Monitoring Instrument (TROPOMI) on board the Sentinel-5 Precursor mission are pivotal in detecting methane emissions (Schneising et al., 2022; Jacob et al., 2022). These technologies provide extensive spatial coverage and detailed data on methane concentrations, which are crucial for identifying emission hotspots and tracking emission trends over time (Sánchez-García et al., 2021).

In addition to the technology used for emission monitoring, several mitigation strategies have been implemented in the dairy industry to reduce GHG emissions. These strategies include dietary modifications to reduce enteric fermentation, manure management techniques such as anaerobic digestion and composting, and precision agriculture practices to minimize over-application of fertilizers (Ghosh et al., (2020); Wattiaux et al., (2019)). Moreover, benchmarking frameworks and decision-support tools like the Integrated Farm System Model (IFSM) and the Cool Farm Tool have been developed to help dairy producers assess the environmental impacts of their operations, identify emission hotspots, and devise mitigation solutions (Arulnathan et al., (2020); Poulopoulou et al., (2023)).

This study harnesses satellite imagery exclusively to assess methane emissions from over 1,000 dairy farms across a five-year period (2019 to 2023), eliminating the inconsistencies associated with ground-level data. This approach provides a clearer and more accurate depiction of emissions across the Canadian dairy landscape, offering a novel perspective on the environmental impact of this sector.

The goal of this research is to develop an AI-driven benchmarking tool that aids in reducing emissions within the Canadian dairy industry. By leveraging advanced analytics and machine learning, this tool is designed to provide stakeholders with actionable insights, enabling the implementation of effective emission reduction strategies and the adoption of sustainable farming practices. The need for such innovations has become increasingly urgent as global warming escalates and as there is a growing push for sustainability and environmental responsibility in agriculture.

Our methodology extends beyond mere data analysis to include the integration and preprocessing of satellite-derived data. This comprehensive framework allows for a holistic view of emission patterns, furnishing dairy farmers with critical insights that can be leveraged for environmental benefits. The study employs a variety of data processing tools, from web scraping for initial data collection to geospatial analysis software and Python scripts for data preprocessing. Subsequently, machine learning algorithms are used for predictive analysis, ensuring each phase of the project is optimized to extract maximum value from the data collected.

The integration of real-time emission data obtained from satellite observations with advanced data analytics enhances the accuracy and efficacy of these tools, providing a more strategic approach to emission reduction. This integration is essential for formulating better recommendations and for making informed decisions aimed at reducing the environmental footprint of dairy farming. The imperative to reduce GHG emissions in the dairy sector is underscored by the need to address global warming and environmental degradation. This study, through its innovative use of satellite technology and AI-driven tools, aims to provide a framework that not only enhances the understanding of emission dynamics but also supports the sustainable development of the dairy industry in Canada.

## Materials and Methods

### Data Collection and Integration

We utilized Octoparse 8, a web scraping tool, to gather information from operational dairy farms and processing facilities across Canada. This extraction focused on ensuring the inclusion of only active establishments in our dataset (Figure S1). The tool enabled us to compile a comprehensive list of dairy farms and processors, forming the foundation for our subsequent analyses. After identifying the locations, we extracted their latitude and longitude coordinates using the Geocode Extension, critical for precise geospatial analysis. These coordinates were organized in Microsoft Excel, sorted by Canadian province, to facilitate further processing and integration. We then converted these Excel sheets into CSV files to comply with the analytical tool standards. These CSV files were uploaded into Google Earth Pro to create KML/KMZ files that visualized the geographic distribution (Figure S2) of these entities. Additionally, we integrated satellite data from Sentinel 5P accessed via Google Earth Engine to enhance our dataset with GHG emission data (Figure S3), enabling us to identify emission patterns and trends across different provinces over time.

### Data Processing and Analysis

The raw data underwent transformation into actionable information through advanced analytics. Initially, data from satellite sources were aggregated and structured using Python scripts, organizing the methane concentration data collected over various weeks. This preprocessing step was essential for organizing and preparing the data for detailed analysis. We employed Power BI for an in-depth analysis of the emissions data, utilizing its robust capabilities to generate graphs and charts that provided insights into emission trends across provinces. Our main objective was to discern patterns in emission rates through exploratory data analysis (EDA), employing time series analysis techniques such as decomposition and forecasting. Furthermore, we applied predictive modeling techniques including autoregressive integrated moving average (ARIMA), Long short-term memory (LSTM), Gradient Boosting Regression Machines (GBR), and PROPHET to develop models capable of forecasting emissions for the next five years. These models, trained on historical data, helped determine the most accurate forecasts for future emissions.

### Emissions Profiling and Trend Analysis

To ensure consistency across the dataset, we conducted thorough data preprocessing, which included cleaning and structuring the emissions data by address. This facilitated a better understanding of emission patterns. We employed time series analysis techniques to decompose the data into components of trend, seasonality, and random fluctuations, aiding in the detailed examination of emissions patterns over time. Statistical methods like the ARIMA were used to predict emission levels from historical data, capable of capturing the complex dependencies within temporal data. Additionally, predictive modeling through machine learning models such as LSTM networks and GBR was utilized to project future emission levels and identify factors influencing emissions. These analyses helped refine our techniques and enhance the accuracy of our predictive models.

### Predictive Analytics for Emission Reduction

We evaluated various models to predict emissions based on historical data, ultimately selecting four; ARIMA, LSTM networks, GBR, and PROPHET for detailed analysis. The development of these predictive models involved training with of historical emissions data. This process enabled the models to learn patterns and relationships within the data. The effectiveness of these models was assessed by calculating the Root Mean Squared Error (RMSE), allowing us to identify the most accurate model for emission forecasting. Further optimizations, such as normalization and data augmentation, were explored to enhance LSTM performance, providing even more precise results in our predictive analytics.

## Results and Discussions Exploratory

### Data Analytics

The data analytics process began with Exploratory Data Analysis (EDA), which facilitated a deeper understanding of the dataset. This phase was crucial for identifying patterns, trends, and anomalies such as outliers or missing values, guiding further data preprocessing. Geospatial analysis played a pivotal role in our project, enabling the visualization of the spatial distribution of dairy farms and processors across various Canadian provinces. Utilizing geographic information system (GIS) tools, including Google Earth Pro and Google Earth Engine, we generated maps and overlays that pinpointed hotspots of greenhouse gas emissions. This analysis was instrumental in forming insights that could assist stakeholders in making informed decisions to mitigate emissions, particularly in identified critical areas. An integral part of our analytical approach involved leveraging Power BI for in-depth analysis and visualization of emission data. Through Power BI, we developed interactive dashboards that provided comprehensive reports, enabling stakeholders to observe and interpret the data effectively. These dashboards included visually engaging charts, graphs, and maps, offering stakeholders valuable insights and highlighting relationships among various factors influencing emissions. The following results were obtained using Power BI.

Utilizing PowerBI, we generated interactive graphs with filter options to analyze emissions by farm, processor, or province, enhancing our understanding of emission distribution and trends.

Methane measurements, recorded by the Sentinel 5P satellite in parts per billion (ppb), indicated that farms significantly outpaced processors in total methane production, with figures reaching 98,500,312.98 ppb for farms compared to 35,985,098.98 ppb for processors (Figure 4 and 5). Ontario emerged as a principal contributor, accounting for 15.51% of the national total, with methane emissions in this province showing a marked increase from 3.4 million ppb in 2019 to 10.5 million ppb by 2023 (Figures 6,7,8 and 9). This rise correlates with the growing number of dairy operations driven by increased demand and the $127 million allocated to enhancing dairy processing infrastructure in Ontario (Dairy Farmers of Ontario 2023 Annual Report).

**Figure 1.**
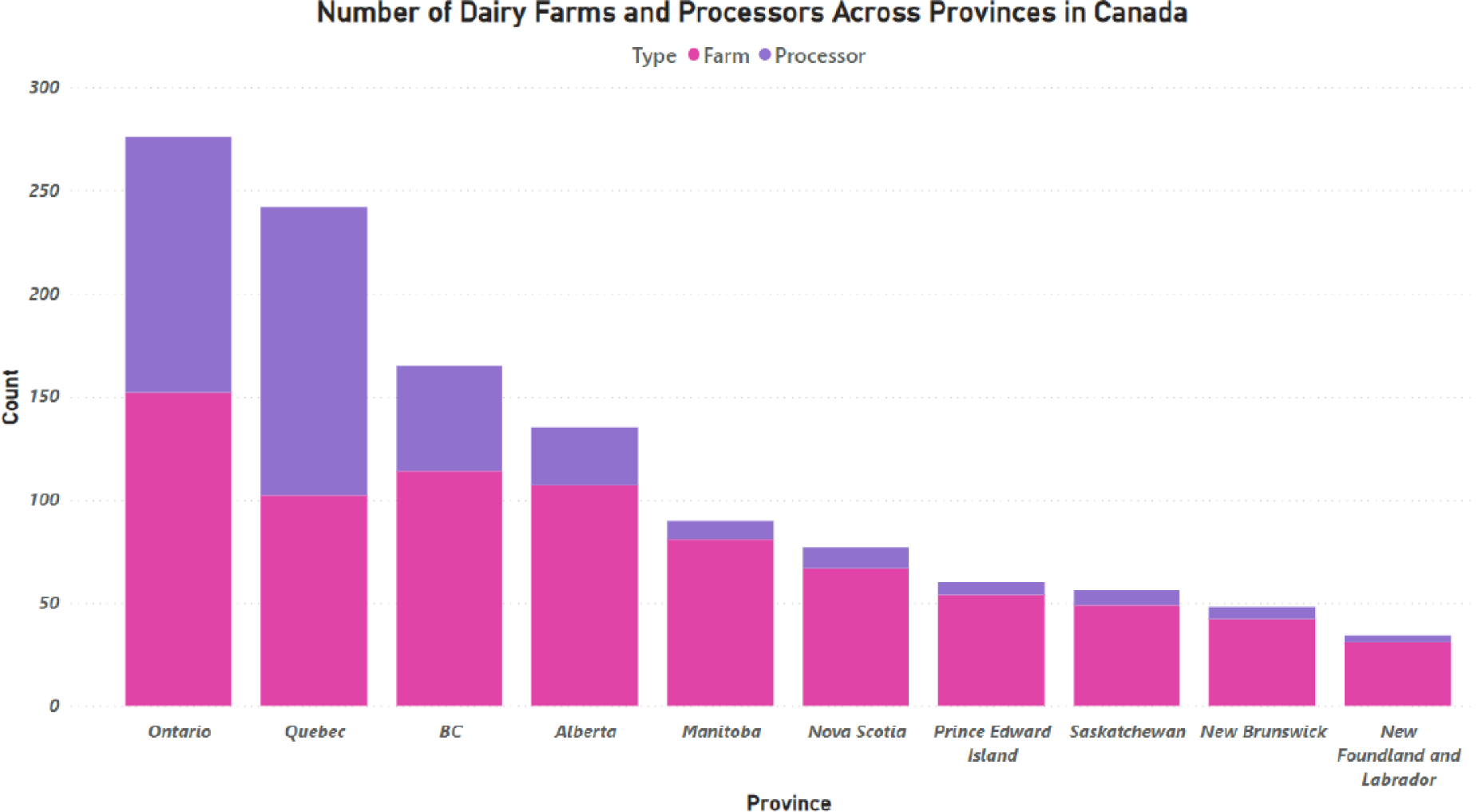
Distribution of Dairy Farms and Processors by Province in Canada.

**Figure 2.**
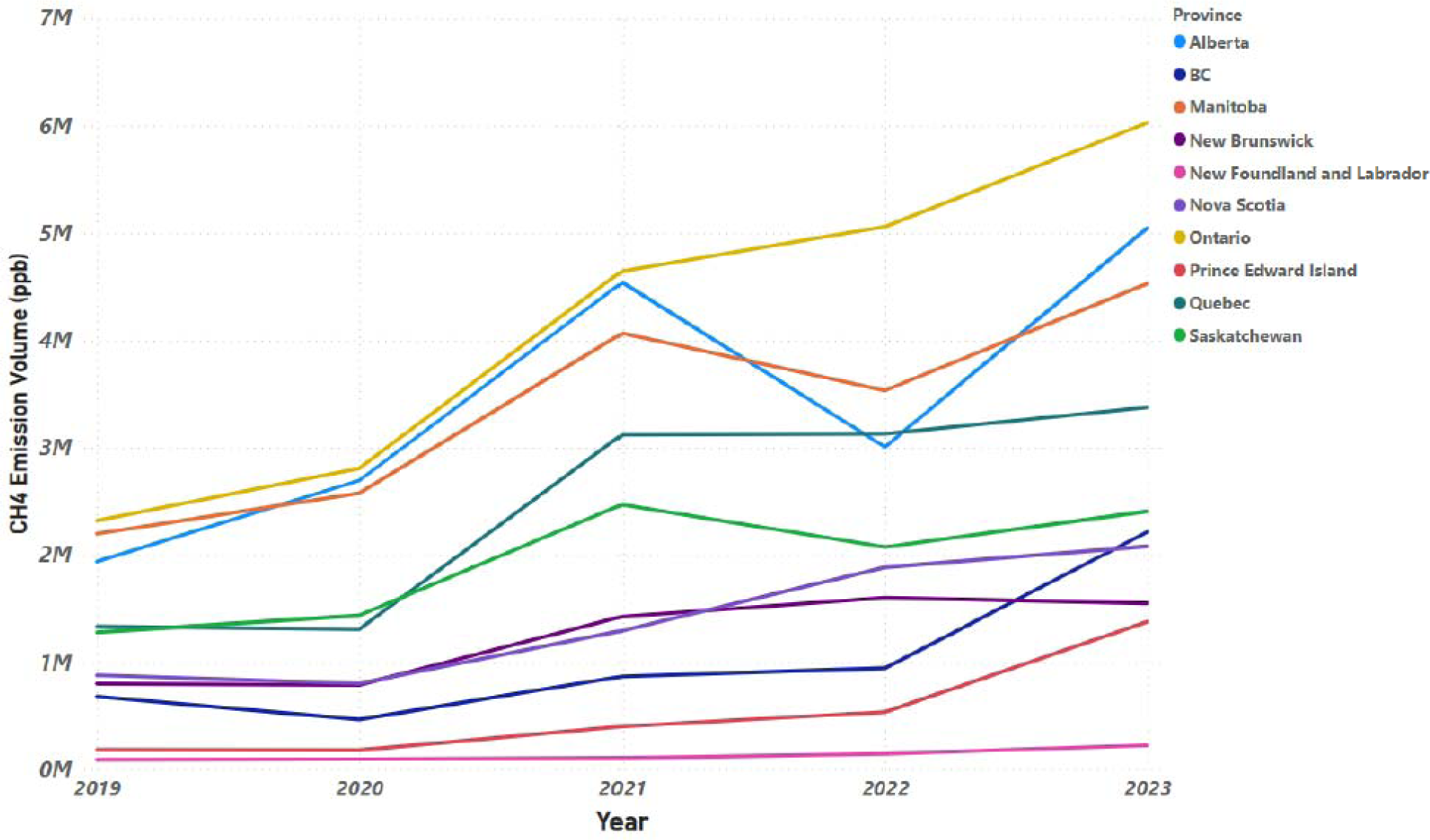
Provincial Methane Emissions from Dairy Farms in Canada (2019-2023).

**Figure 3.**
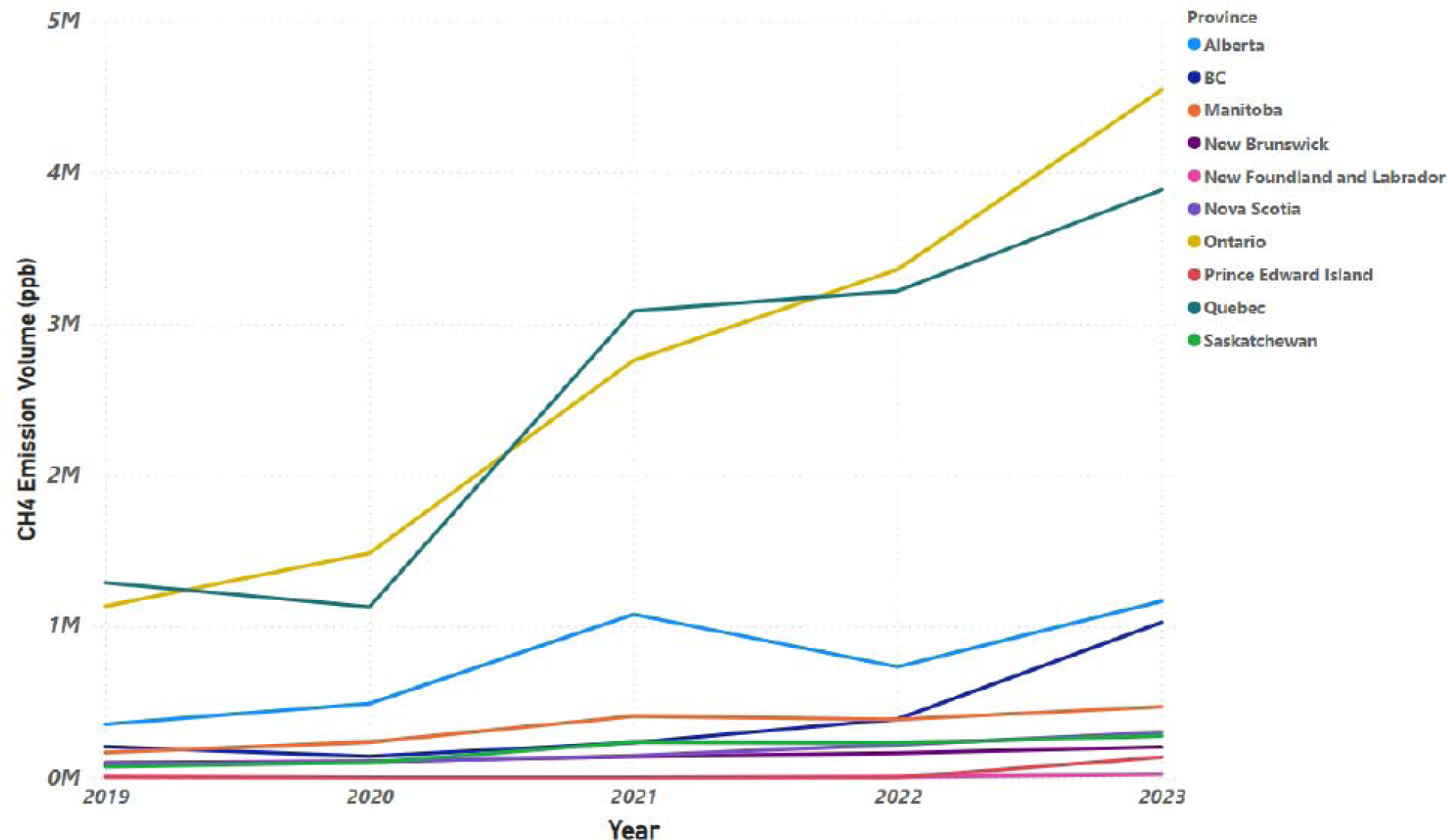
Provincial Methane Emissions from Dairy Processors in Canada (2019-2023).

**Figure (4A).**
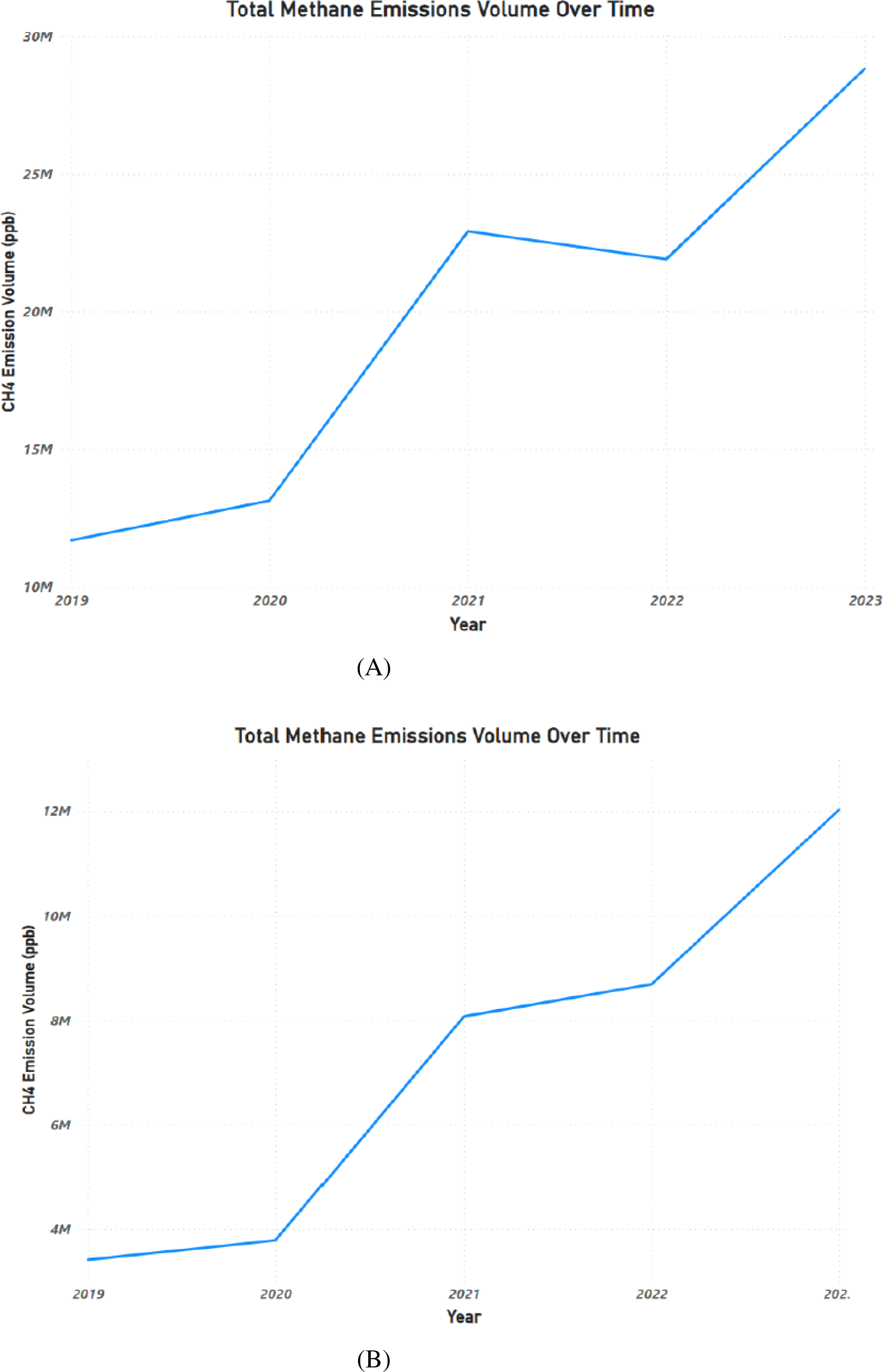
Total Methane Emissions from Canadian Dairy Farms. (4B). Total Methane Emissions from Canadian Dairy Processors.

**Figure 5.**
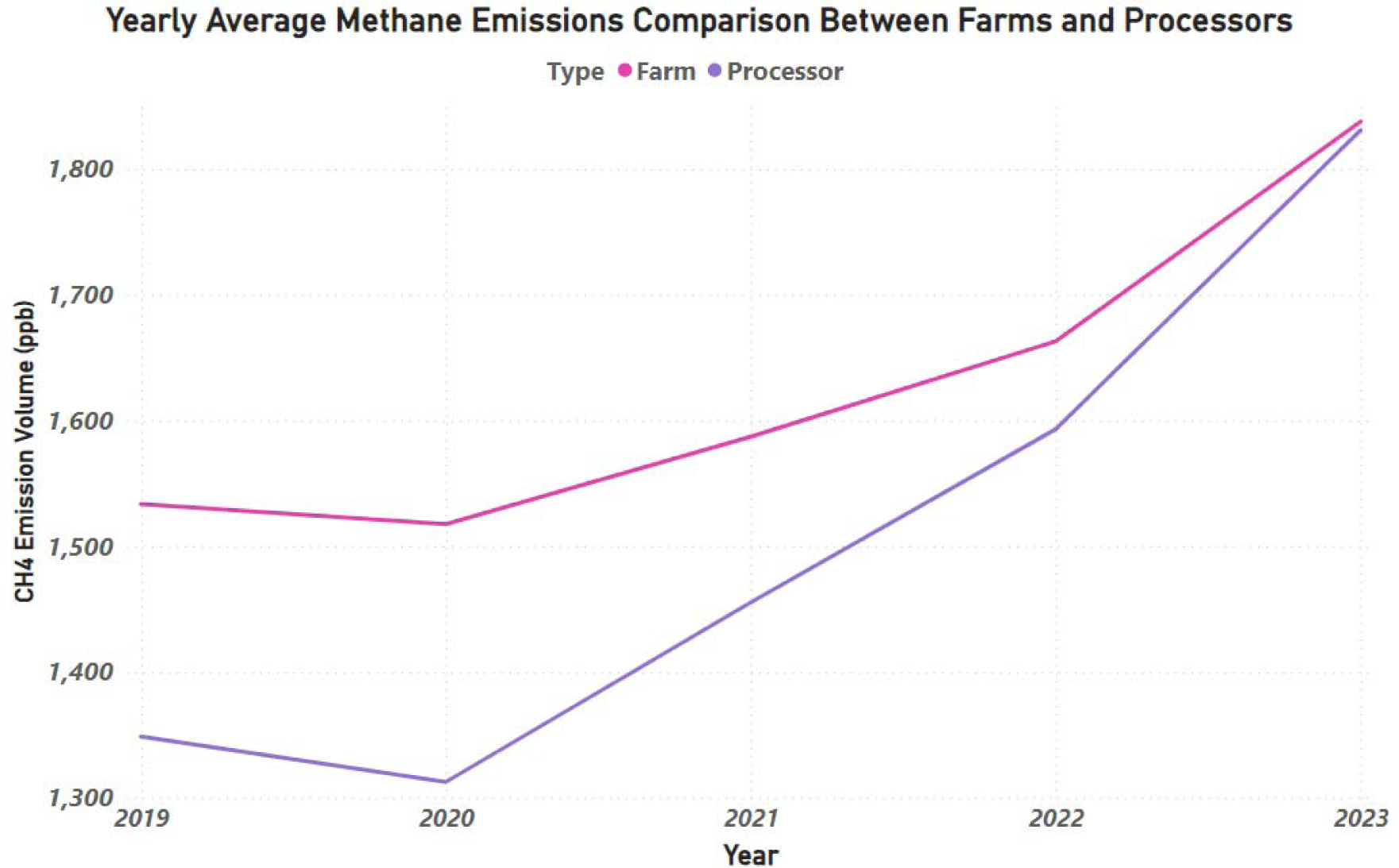
Annual Average Methane Emissions from Canadian Dairy Processors (2019-2023).

**Figure 6.**
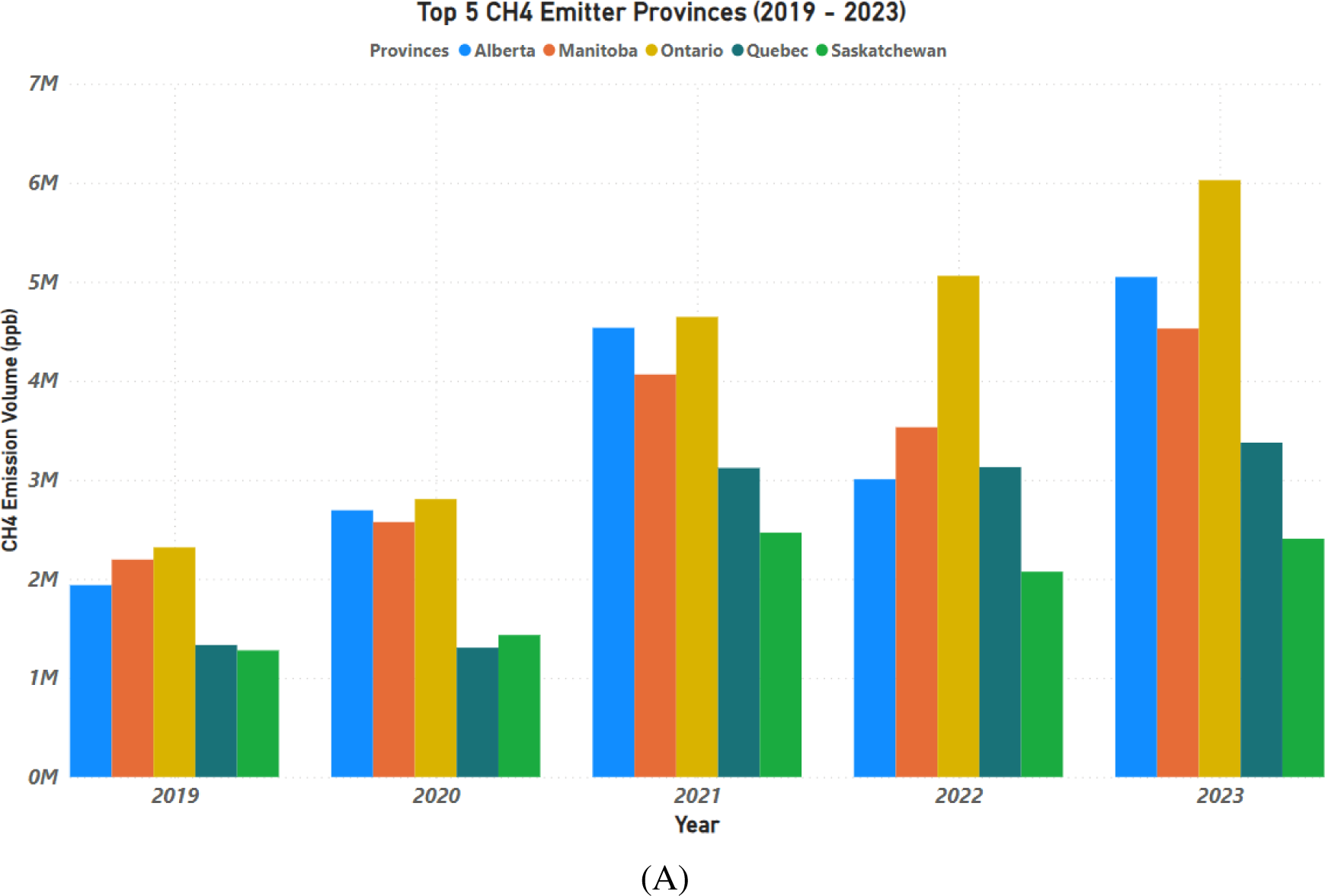
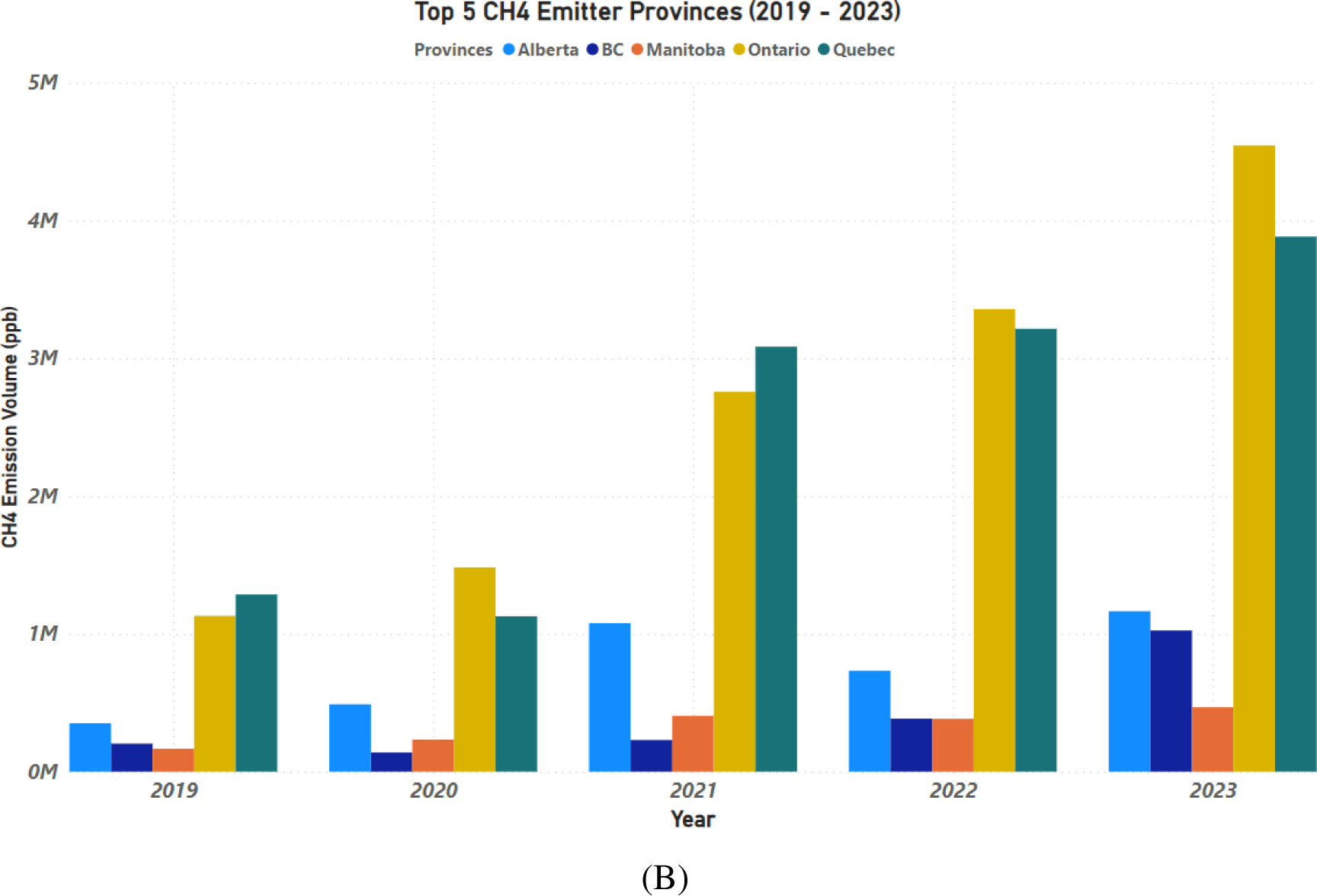
Canadian provinces with the Highest Methane Emissions from Dairy Farms (A) and Dairy Processors (B) between the year 2019 to 2023.

**Figure 7.**
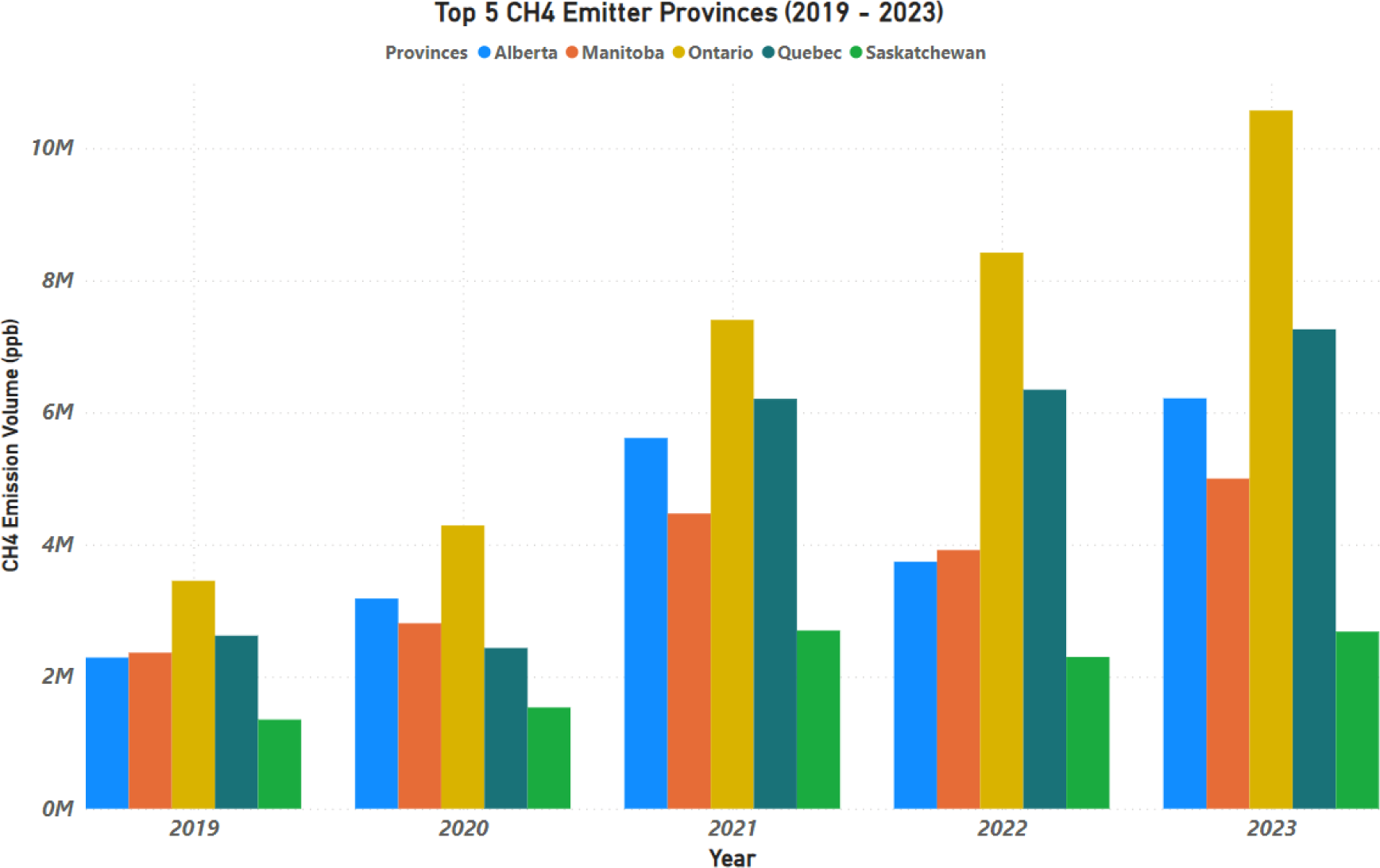
Top Methane Emitting Provinces in the Canadian Dairy Sector.

**Figure 8.**
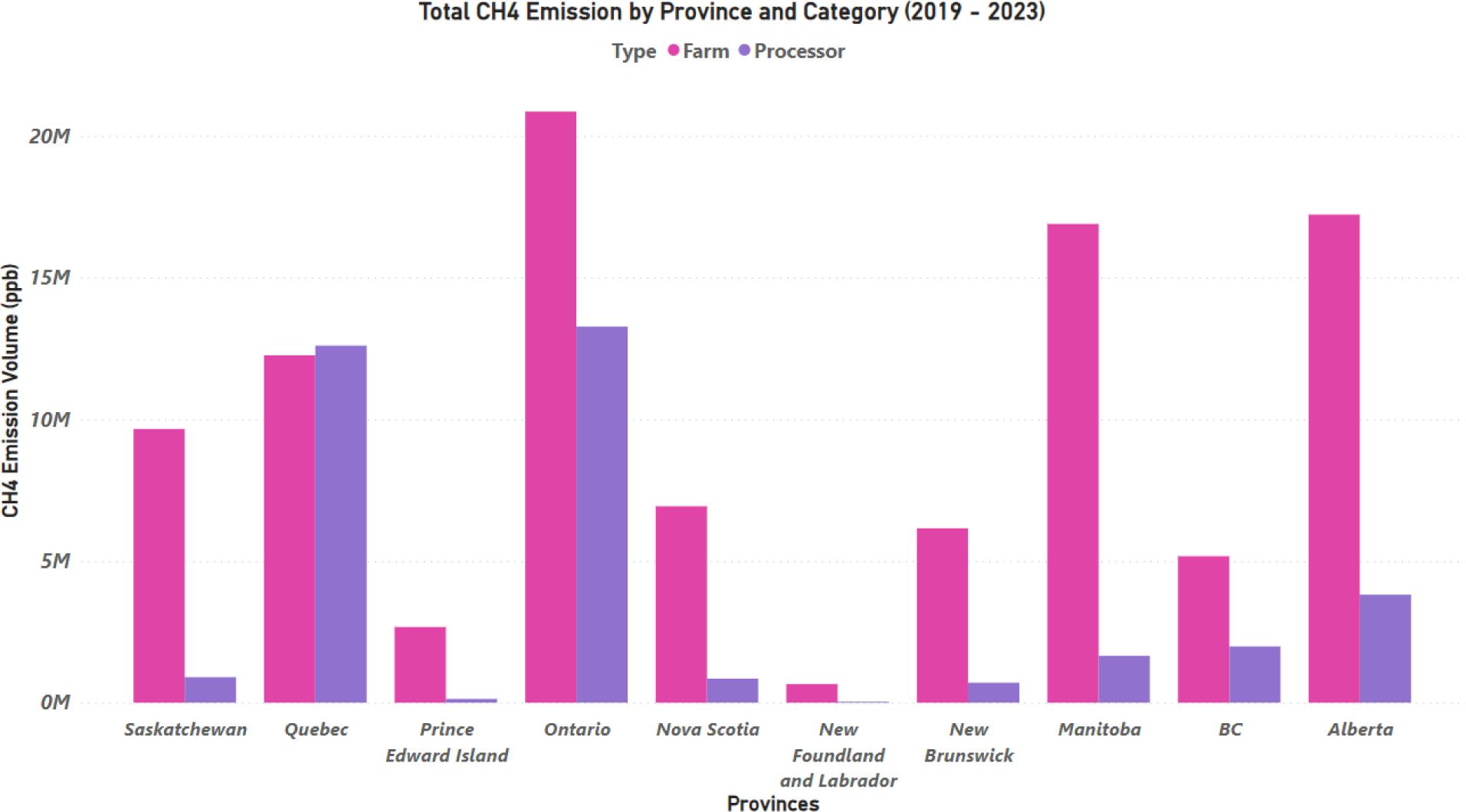
Provincial Total Methane Emissions from Dairy Farms and Processors between the year 2019 to 2023.

Nationwide, methane emissions escalated from 15.1 million ppb in 2019 to approximately 40.8 million ppb in 2023, with farms contributing nearly 60% of the total in 2023 (Figure 9). A notable reduction in emissions was observed in 2022 across several provinces, attributed to the pandemic’s impact on the dairy industry, leading to decreased cow populations and, consequently, reduced methane output (Figures 6 and 9).

**Figure 9A.**
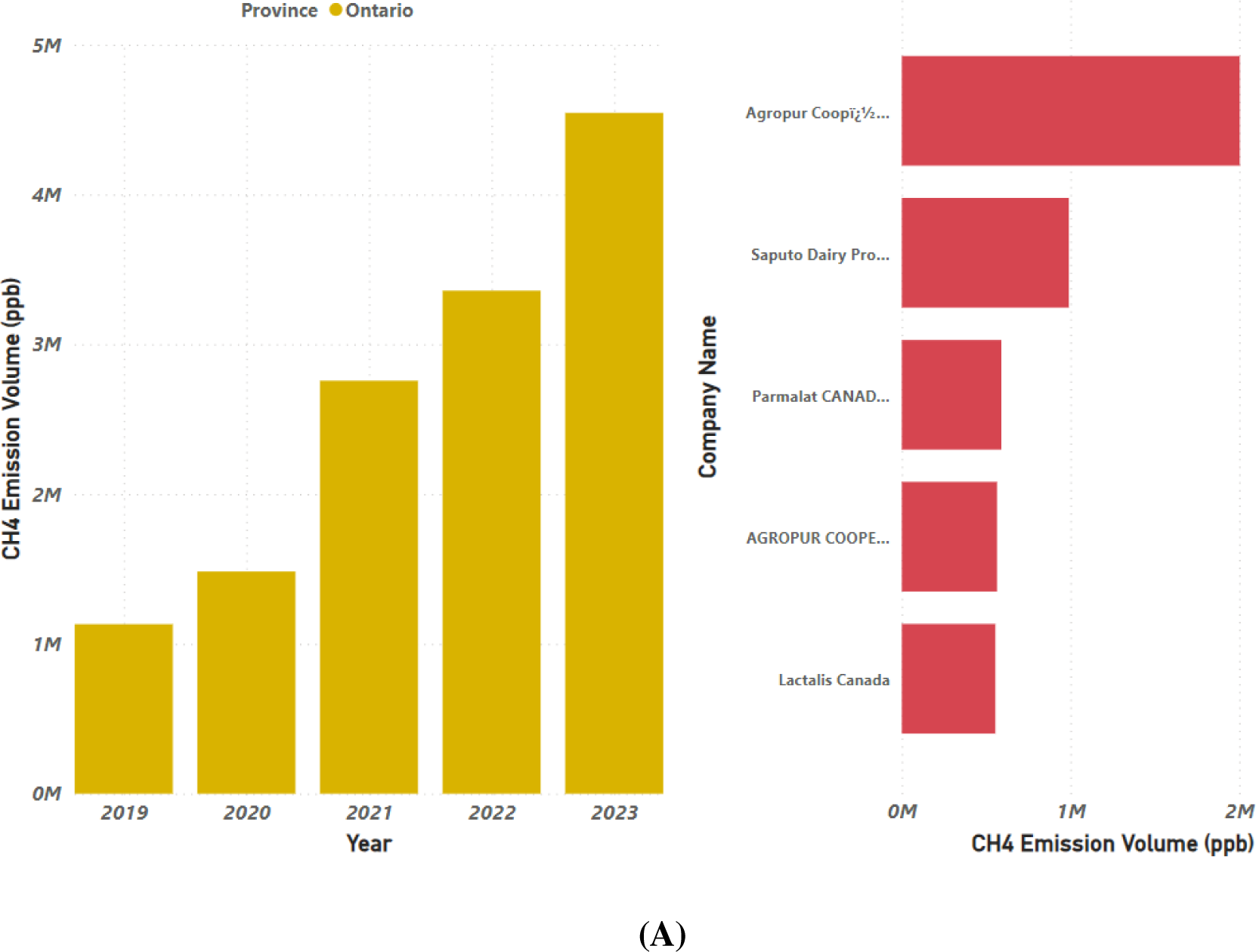
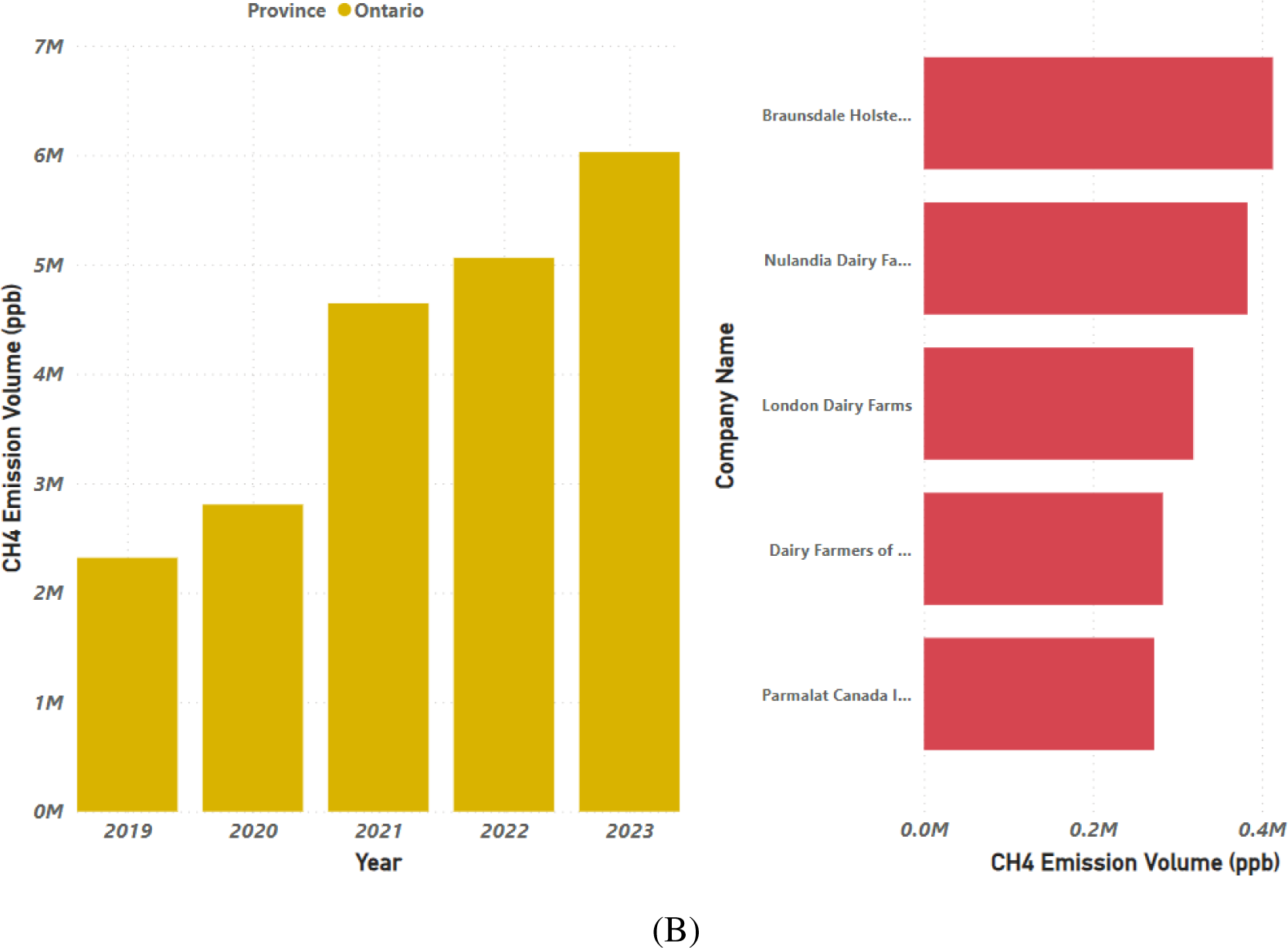
Highest Annual Methane Emissions by Province and Top Emitting Dairy Processors in Canada (A). Highest Annual Methane Emissions by Province and Top Emitting Dairy Producers (B) in Canada.

Our predictive modeling utilized advanced machine learning algorithms to forecast future emissions and guide emission reduction strategies effectively. We assessed several models,

ARIMA (Autoregressive Integrated Moving Average): This model excels in capturing data trends through its combined autoregressive and moving average components.

PROPHET: Developed by Facebook, this additive regression model is tailored for seasonal effects and recognizes holiday impacts in time series data.

Gradient Boosting Regression (GBR): An ensemble learning method that enhances predictions by integrating multiple weak decision trees to correct previous errors incrementally.

Long Short-Term Memory (LSTM) networks: Ideal for sequential data, LSTMs manage long-term dependencies within time series, making them highly effective for this analysis.

These models, trained on historical data, learned to identify intricate patterns and relationships, providing accurate emission forecasts. LSTM demonstrated superior precision, reflected by its minimal Root Mean Square Error (RMSE), although further attempts to enhance its accuracy showed no improvement in RMSE values.

### Assessing Methane Emissions in Canadian Dairy Farms Through AI-Driven Analytics

The development of an AI-driven benchmarking tool designed for emission reduction across Canadian dairy farms and processors has yielded significant insights and actionable recommendations for the industry. Through an integrated approach involving data collection, integration, preprocessing, and predictive analysis, our study has identified key patterns, trends, and predictive models aimed at mitigating emissions within the dairy supply chain.

### Geographical and Temporal Emission Patterns

One of the pivotal findings of our study was the geographical variance in methane emissions across Canadian provinces. Geospatial analysis and visualization techniques pinpointed emission hotspots, with Alberta, Ontario, and Quebec consistently ranking as the top three emitters for both dairy farms and processors. For example, in 2019, Alberta contributed 28% of the total methane emissions from Canadian dairy farms, followed by Ontario with 25%, and Quebec with 20%. These data have enabled stakeholders to focus mitigation efforts on these high-emission provinces. Temporal analysis of the emissions data revealed seasonal patterns, with peak emissions occurring during the summer months. This seasonal increase is likely due to heightened microbial activity in warmer temperatures, which accelerates methane production. Over the five-year span from 2019 to 2023, there was an observed overall increase in methane emissions by approximately 7.2%.

### Predictive Modeling and Emission Forecasting

Our evaluation of various machine learning models identified the LSTM network as the most effective, demonstrating the lowest RMSE value of 15.40 for predicting future emissions. The LSTM model’s ability to capture complex temporal dependencies in the emission data underscores its utility for ongoing monitoring and forecasting. The choice of RMSE as the primary metric for model evaluation was strategic, given its sensitivity to large errors—a crucial feature that ensures high-stakes prediction errors are adequately penalized, enhancing the reliability of emission forecasts. However, RMSE’s sensitivity to data anomalies can sometimes skew analysis, necessitating careful handling of outlier data to maintain accuracy.

### Feature Importance and Emission Influencers

Feature importance analysis shed light on the primary drivers of emissions in dairy operations. Key factors such as feed composition, manure management, and herd size were identified as significant contributors to methane emissions. Notably, a 10% increase in herd size correlated with a 6.8% rise in methane emissions. Moreover, regional climatic conditions also emerged as critical determinants of methane output, emphasizing the need for region-specific mitigation strategies.

### Impact of Dairy Herd Size and Production on Emissions

The analysis highlighted a direct correlation between herd size and methane emissions. Farms with over 500 cows were responsible for 60% of total methane emissions from Canadian dairy operations. This finding illustrates the substantial environmental impact of large-scale dairy farms and underscores the importance of scalable emission reduction strategies. Additionally, a strong link was observed between milk production volumes and methane emissions. Increased milk production, typically associated with larger herd sizes and more intensive enteric fermentation processes, directly influences methane output. For instance, for every 1,000 liters of milk produced, approximately 22 kg of methane was emitted (Figure 11). This relationship highlights the environmental costs of high-production dairy farming and the necessity for interventions that enhance feed efficiency, optimize herd management, and incorporate advanced emission-reducing technologies.

**Figure 10.**
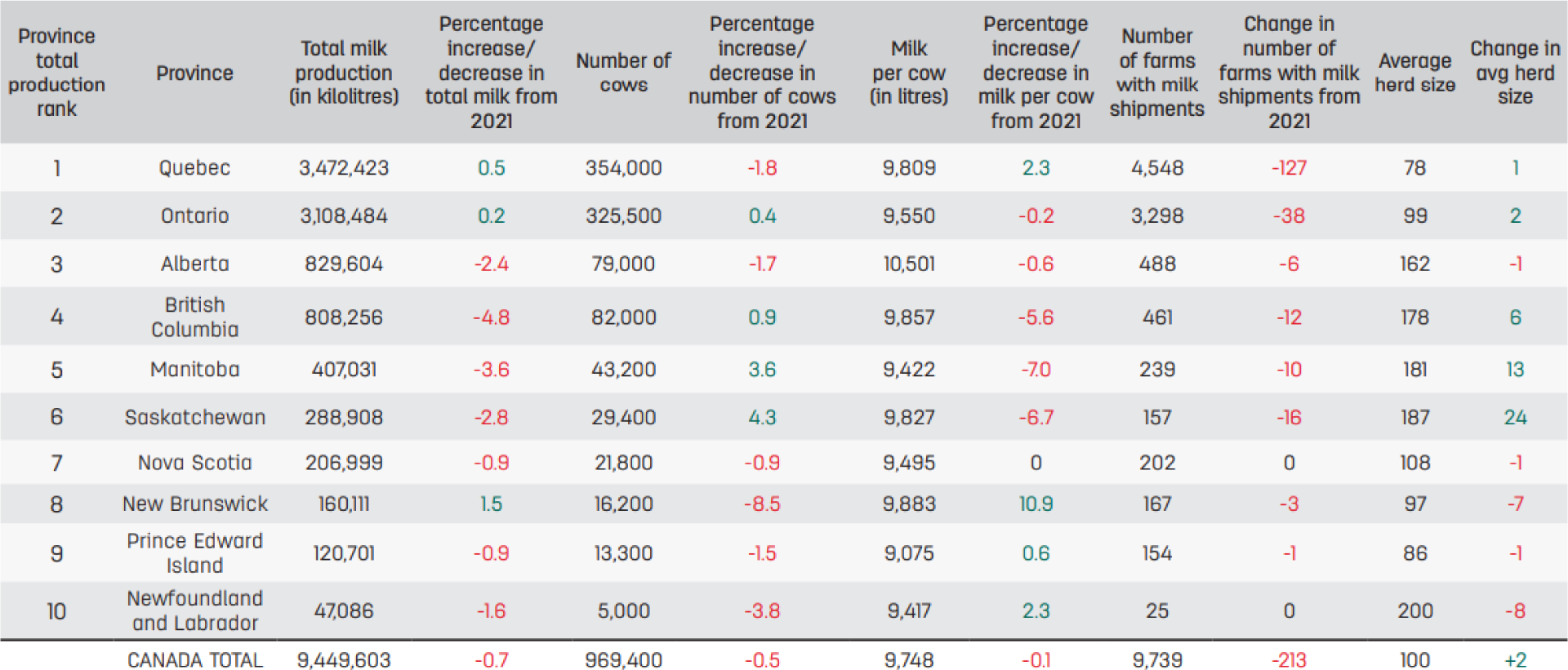
Dairy Production Statistics for 2022 Across Canadian Provinces - Milk Production and Cattle Numbers. (Progressive Dairy, 2022)

**Figure 11.**
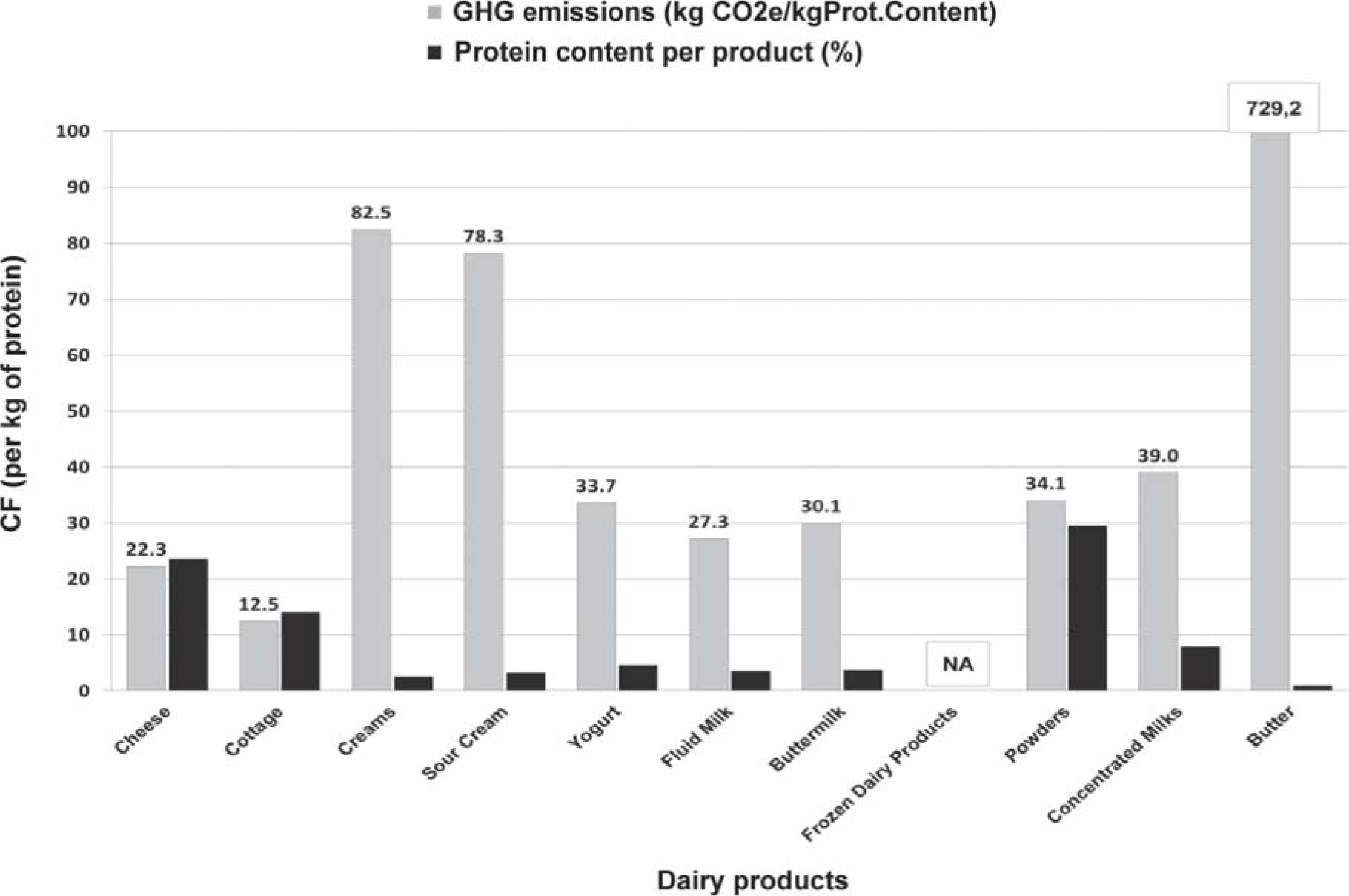
Carbon Footprint of Canadian Dairy Products per Kilogram of Protein Content. This figure illustrates the greenhouse gas (GHG) emissions in CO_2_ equivalents (CO_2_e) for various dairy products (Verge et al., 2013).

**Figure 12.**
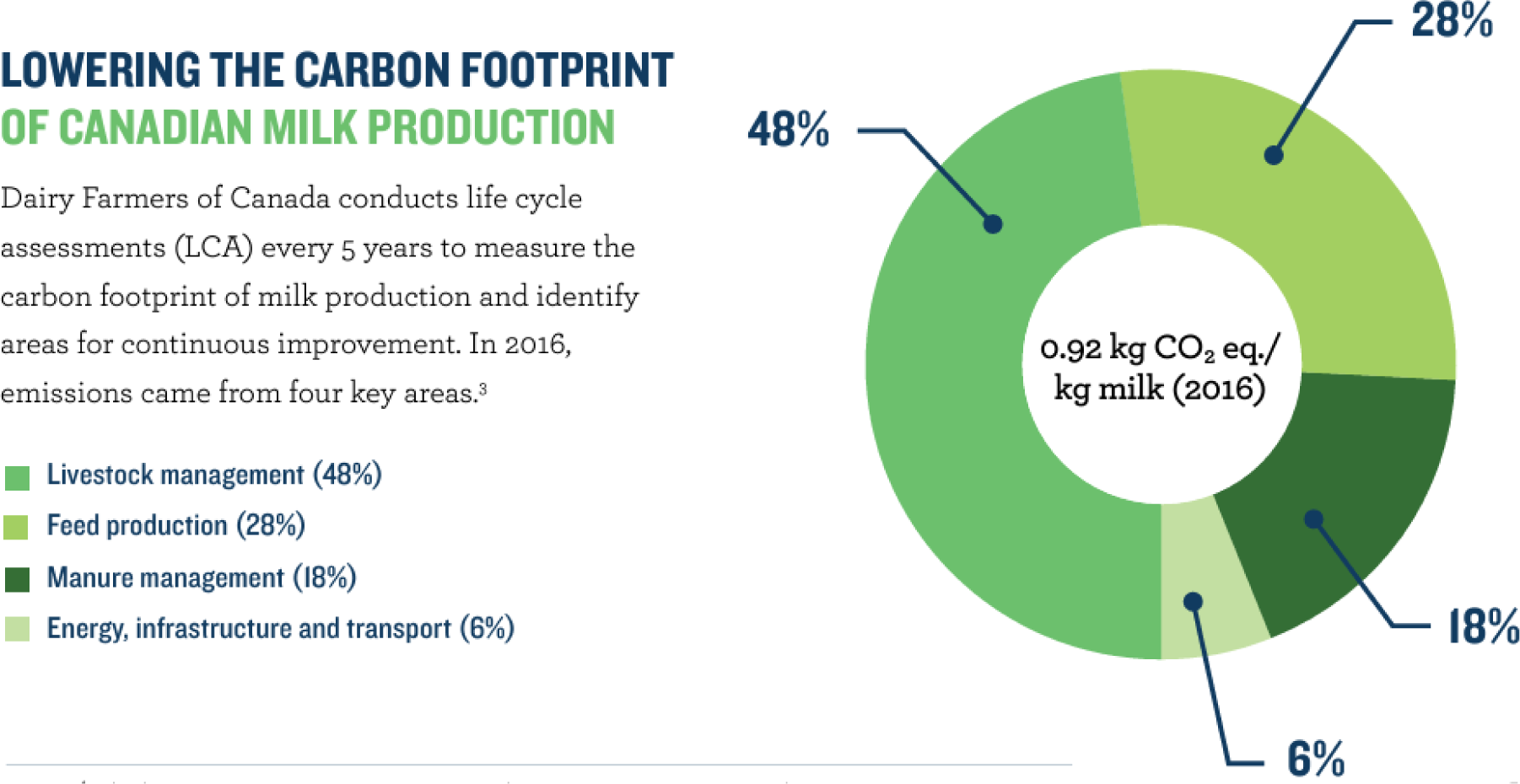
Distribution of Emission Sources in Canadian Milk Production. Pie chart illustrating the primary sources of emissions in Canadian dairy farming as reported in [Dairy Farming Forward to 2050 Report, 2023]. The chart details percentages from livestock management (48%), feed production (28%), manure management (18%), and combined energy, infrastructure, and transport (6%).

### Strategies and Performance in Emission Reduction in the Canadian Dairy Industry

Over the period from 2019 to 2023, the Canadian dairy industry exhibited a modest average annual increase in emissions of 1.2%, suggesting some effectiveness in existing mitigation strategies and improved farm management practices. However, reversing the overall upward trend and achieving significant long-term reductions require more robust and sustained interventions (Environment and Climate Change Canada, 2023).

Despite its progress in reducing GHG emissions and achieving a relatively low global carbon footprint for milk production, the Canadian dairy industry still has substantial room for improvement, particularly within the dairy processing sector. The sector accounts for less than 1.3% of Canada’s total GHG emissions, with primary sources being enteric fermentation (48%), feed production (28%), and manure management (18%) (Neethirajan, 2023).

### Addressing Emissions from Dairy Processors

Emissions from dairy processors, though less significant than those from farms, still contribute notably to the industry’s carbon footprint. Key areas include energy use in processing activities such as pasteurization, drying, and emissions linked to transportation of processed products and packaging materials. While specific data on emissions from processors are scarce, estimates suggest that these areas together account for about 6% of the industry’s total emissions (Neethirajan, 2023). Strategic improvements in production efficiency, adoption of alternative energy sources, sustainable packaging solutions, and enhanced waste management could significantly reduce emissions from this segment. Such initiatives are critical for the dairy industry to further decrease its overall carbon footprint.

### Evaluation of Predictive Models for Emission Forecasting

In our comparative analysis of predictive models, the Long Short-Term Memory (LSTM) model demonstrated superior performance, with the lowest Root Mean Square Error (RMSE) of 15.40, indicating its proficiency in capturing complex nonlinear patterns and long-term dependencies in emission data. This makes it particularly useful for reliable forecasting in the context of emissions management. Although the LSTM model offers limited interpretability compared to other models like ARIMA and GBR, its accuracy in forecasting emissions is crucial for strategic planning. The GBR and ARIMA models, which scored RMSEs of 19.53 and 22.63 respectively, showed capability in handling nonlinear and linear patterns but did not perform as well as LSTM. The Prophet model lagged behind with an RMSE of 37.22, reflecting its inadequate suitability for this specific application.

### Leveraging Data Insights for Targeted Emission Reduction

The insights obtained from our comprehensive data analytics enable stakeholders to implement more targeted and effective mitigation measures. Regions with high emissions intensity, such as provinces identified through our analysis, can focus on investments in advanced manure management systems, precision agriculture techniques, and the promotion of best practices among large dairy operations. Additionally, dairy processors could concentrate efforts on improving energy efficiency, exploring sustainable packaging options, and optimizing transportation logistics to further reduce their emissions impact.

### Economic Contributions and Environmental Initiatives of the Canadian Dairy Industry

The Dairy Farmers of Canada and the Dairy Processors Association of Canada provide critical economic and environmental insights. The dairy sector generated $7.4 billion in farm cash receipts and $16.2 billion from dairy product sales in 2021, marking significant contributions to the national economy (Neethirajan, 2024; Canadian Dairy Commission Annual Report 2022).

In the year 2021, Canada hosted 9,952 dairy farms with a total dairy cow population of 981,300, producing 396 million kg of butterfat. Ontario is a major contributor, with over 3 billion liters of milk produced annually. Canada is known for its diverse range of over 1,450 cheese varieties and maintains stringent quality control standards, enhancing its global reputation. While there was a slight decrease in the consumption of certain dairy products in 2021-2022, there was an uptick in the consumption of higher-fat products like cream and butter, reflecting changing consumer preferences. Notable initiatives include the Pathways to Dairy Net Zero, aiming to enhance production efficiencies and reduce emissions, exemplifying the industry’s commitment to environmental sustainability (Dairy Farmers of Ontario 2023 Annual Report).

The results from our study underscore the Canadian dairy industry’s dedication to balancing economic growth with environmental stewardship, setting a benchmark for global dairy standards. By harnessing advanced analytics and predictive modeling, stakeholders are better equipped to formulate and execute effective strategies that align with Canada’s broader climate goals, demonstrating leadership in sustainable dairy production and processing practices.

### Scope 1, 2, and 3 Emissions in the Dairy Industry

In the context of greenhouse gas accounting and emissions reduction in the dairy industry, it’s essential to understand the classification of emissions into Scope 1, Scope 2, and Scope 3. These scopes are defined by the Greenhouse Gas Protocol, which provides the world’s most widely used greenhouse gas accounting standards (Kuramochi et al., 2020).

Scope 1 Emissions are direct emissions from owned or controlled sources. In the dairy industry, these emissions primarily come from enteric fermentation in ruminants—where methane is produced by cows as part of their digestive process—and from manure management systems that release methane when manure is stored or treated anaerobically. Scope 1 also includes emissions from fuels burned on farms for heating or in farm vehicles.

Scope 2 Emissions are indirect emissions from the generation of purchased electricity, steam, heating, and cooling consumed by the reporting company. For dairy farms and processors, Scope 2 emissions arise from the use of electrical power to operate milking machines, cooling systems, and other processing equipment such as pasteurizers and dryers.

Scope 3 Emissions are all indirect emissions (not included in Scope 2) that occur in the value chain of the reporting company, including both upstream and downstream emissions. In the dairy industry, these include emissions associated with the production of purchased goods, such as feed and veterinary drugs; transportation services; emissions from the production of fuels and energy not covered under Scope 2; and emissions resulting from the use of sold products. For processors, this includes emissions linked to transportation of processed products, packaging production, and waste disposal.

### Integrating Scope Emissions into Reduction Strategies

Understanding these emissions categories helps the dairy industry identify and prioritize areas for emission reduction. It aligns with strategies to reduce Scope 1 emissions through improved manure management techniques and dietary changes to reduce enteric fermentation. Technologies such as anaerobic digesters can capture methane from manure and convert it to energy, directly reducing Scope 1 emissions while indirectly cutting Scope 2 emissions by decreasing the need for purchased energy. Improvements in energy efficiency and transitions to renewable energy sources like solar or wind can substantially lower Scope 2 emissions. For instance, installing solar panels at dairy processing facilities can reduce reliance on grid electricity, which is typically generated from fossil fuels. Addressing Scope 3 emissions involves engaging with suppliers to ensure that feed is produced with lower emissions and optimizing logistics to reduce emissions from transportation. It also includes working with packaging suppliers to find sustainable options that require less energy to produce and can be recycled or are biodegradable, reducing waste management emissions.

### Broader Implications and Strategic Approaches

By analyzing Scope 1, 2, and 3 emissions, dairy industry stakeholders can develop a comprehensive view of their overall carbon footprint. This holistic approach is crucial for setting realistic emission reduction targets and aligning with global standards such as the Paris Agreement and Sustainable Development Goals. It also helps the industry anticipate regulatory changes and prepare for potential carbon pricing mechanisms, ensuring economic sustainability alongside environmental responsibility. Implementing strategies to reduce these emissions can also enhance the dairy industry’s public image, appealing to environmentally conscious consumers and investors interested in sustainable agricultural practices. Moreover, it can open up new opportunities for funding and partnerships, as many governments and organizations are willing to invest in projects that aim to reduce greenhouse gas emissions and promote sustainability in agriculture.

## Discussions

Our analysis underscores pronounced regional disparities in methane emissions, pinpointing Alberta, Ontario, and Quebec as major contributors. This geographical specificity necessitates tailored mitigation strategies that account for local climatic conditions, farming practices, herd sizes, and the availability of advanced technologies. Targeting these high-emission areas allows for more efficient resource allocation and the implementation of impactful interventions like sophisticated manure management systems, precision agriculture, and the adoption of cutting-edge sustainable farming practices. This approach not only optimizes resource use but also ensures that mitigation strategies are contextually relevant and effective, adhering to the principle that environmental management solutions should be as localized as the challenges they aim to address. The distinct seasonal variations in emissions, with peaks during summer months due to increased microbial activity, suggest the need for dynamic operational strategies. These findings can guide dairy operations in scheduling emission mitigation efforts, such as optimized manure management during high-risk periods. Additionally, adjusting feeding practices to reduce enteric fermentation during warmer months could substantially lower methane outputs.

The deployment of the Long Short-Term Memory model as a predictive tool is an advancement in forecasting emission trends and guiding strategic decisions in the dairy industry. The LSTM’s robust performance in modeling complex patterns enables stakeholders to anticipate future emission scenarios and tailor their strategies accordingly. This predictive prowess is instrumental in transitioning from reactive to proactive environmental management, enhancing the industry’s ability to meet both immediate and long-term sustainability goals.

Our findings also highlight the correlation between increased milk production and elevated methane emissions. This relationship underscores the challenge of balancing productivity with environmental stewardship. Innovative strategies that decouple productivity from emissions—such as improving feed efficiency and optimizing herd management—are crucial. These strategies not only address environmental concerns but also bolster economic efficiency, ensuring the industry’s sustainability in the broadest sense. Benchmarking emissions against global standards and conducting comparative analyses with other leading dairy-producing nations can provide additional layers of insight and drive improvements. These benchmarks can serve as a basis for setting realistic, achievable emissions targets that also align with international best practices, facilitating a global perspective on sustainable dairy production.

Understanding the different scopes of emissions is critical for comprehensive environmental management. Scope 1 emissions include direct emissions from owned or controlled sources, while Scope 2 covers indirect emissions from the generation of purchased electricity, steam, heating, and cooling consumed by the reporting company. Scope 3 emissions, often the most challenging to calculate and reduce, include all other indirect emissions that occur in a company’s value chain. In the context of the dairy industry, these can involve emissions related to feed production, purchased goods and services, transportation and distribution, and waste disposal. Effective management of these emissions requires a holistic approach, integrating detailed tracking, innovative reduction strategies, and consistent stakeholder engagement across the supply chain.

*From the insights gained, several actionable items emerge,*

Targeted Interventions in High-emission Provinces - Implementing specific technological and management interventions in regions identified as high emitters.

Seasonal Strategy Adjustments - Aligning mitigation efforts with seasonal emission peaks to optimize their effectiveness.

Adoption of Advanced Predictive Models - Utilizing LSTM and other advanced models to forecast emissions and inform long-term strategic planning.

Sustainable Practice Implementation - Encouraging widespread adoption of proven sustainable practices through policy support, financial incentives, and educational programs. Continual Improvement through Benchmarking - Regularly updating benchmarks and comparative analyses to keep pace with global best practices and technological advancements.

### Conclusions

The development of an Artificial Intelligence-driven benchmarking tool for emission reduction in Canadian dairy farms marks a significant technological advancement aimed at addressing environmental challenges within the agricultural sector. Our study utilized a comprehensive approach by integrating satellite-derived methane emission data with advanced machine learning techniques, notably geospatial analysis and predictive modeling, to monitor and analyze emissions from over 1,000 dairy farms and processors across Canada. Central to this study was the employment of the Long Short-Term Memory model, which demonstrated superior accuracy in forecasting methane emissions, as evidenced by its minimal Root Mean Squared Error. This model’s ability to capture complex temporal dependencies proved crucial for providing stakeholders with precise, real-time insights into emission dynamics, thereby facilitating informed decision-making and effective implementation of targeted emission reduction strategies.

The investigation revealed significant geographical disparities in methane emissions across the regions of Alberta, Ontario, and Quebec, pinpointing them as major emission hotspots. This finding underscores the necessity for localized mitigation strategies that are specifically tailored to the unique environmental and operational conditions of these regions. Additionally, the study highlighted the critical role of seasonal variations in emissions, with peaks during warmer months due to increased microbial activity. This seasonal pattern suggests that mitigation efforts could be strategically scheduled to maximize their efficacy during periods of elevated emissions. Moreover, the analysis extended beyond farm-level operations to include the dairy processing segment, identifying substantial opportunities for emission reductions through enhanced energy efficiency, sustainable packaging, and optimized waste management. Although dairy processors contribute less to the overall emissions compared to farms, addressing these aspects could lead to significant decreases in the industry’s carbon footprint.

The predictive capabilities of the LSTM model enhance strategic planning within the dairy industry by allowing stakeholders to forecast future emission scenarios and assess the potential impact of various interventions before their implementation. This forward-looking approach is crucial for aligning operational strategies with environmental targets and adapting to potential regulatory changes that could impact the industry. By detailing the interplay between increased milk production, herd size, and methane emissions, the study also stresses the need for innovative strategies that decouple productivity from environmental impact. Such strategies may include improving feed efficiency, optimizing herd management, and incorporating advanced emission-reducing technologies.

## Supporting information

Codes Supplementary FIle

## Data Availability Statement

Anonymized data from this study may be available upon reasonable request.

## Author Contributions

Pratik Parmar: Original draft preparation, writing review and editing, visualization, formal analysis, and validation.

Hangqing Bi: Formal analysis and validation.

Suresh Neethirajan: Conceptualization, supervision, methodology development, and manuscript review.

## Supplementary Files

Supplementary information, including Python codes, PowerBI algorithms, and figures (S1, S2, S3), is available as part of the supplementary files.

## Acknowledgements

The authors thank the Natural Sciences and Engineering Research Council of Canada (NSERC), Mitacs Canada, Net Zero Atlantic Canada Funding Agency, and the New Brunswick Department of Agriculture, Aquaculture and Fisheries for funding this study.

