## Supplementary figures and images for "Artificial Intelligence driven Benchmarking Tool for Emission Reduction in Canadian Dairy Farms"

### Figure S1.png

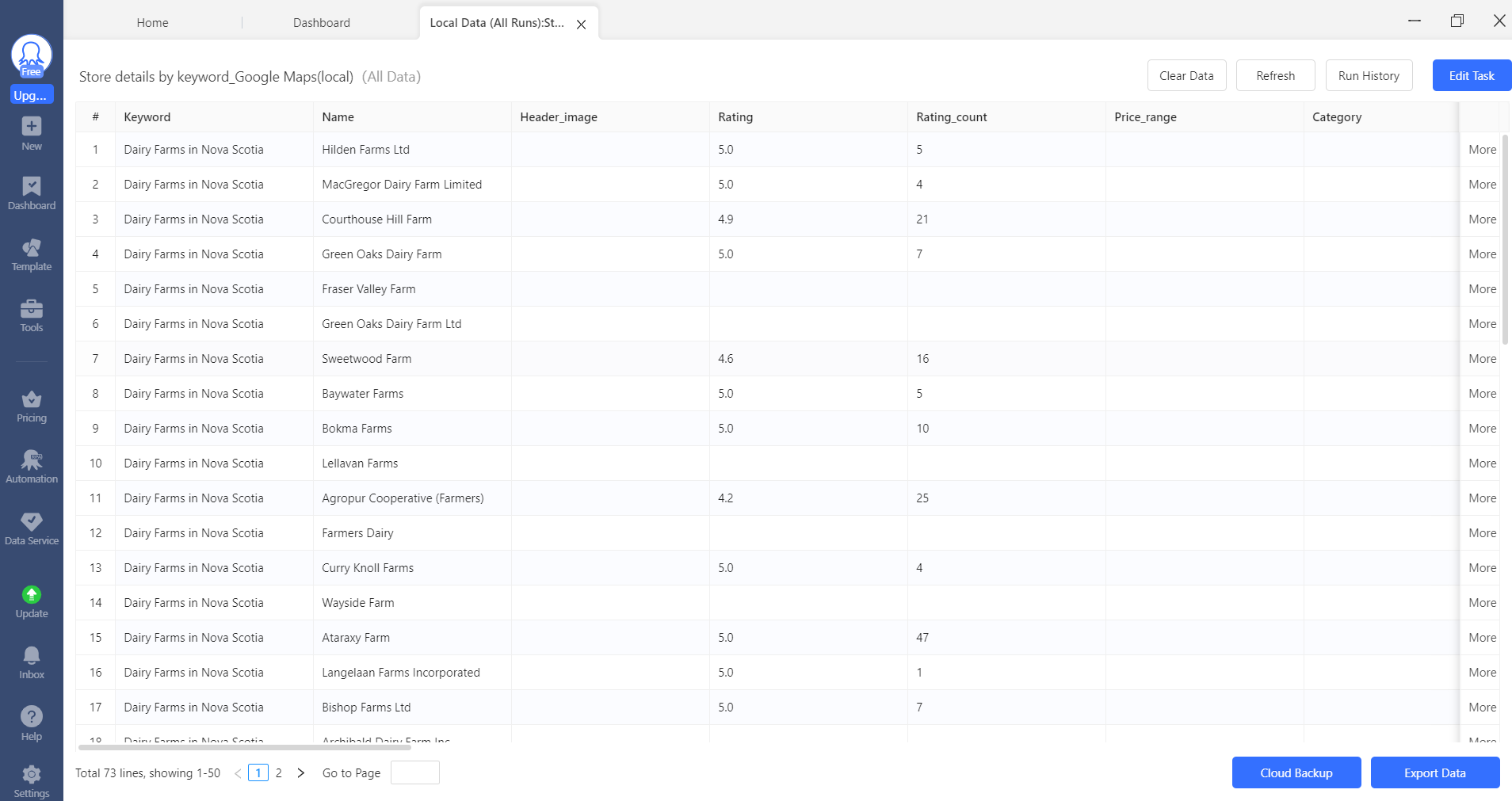

### Figure S2.jpg

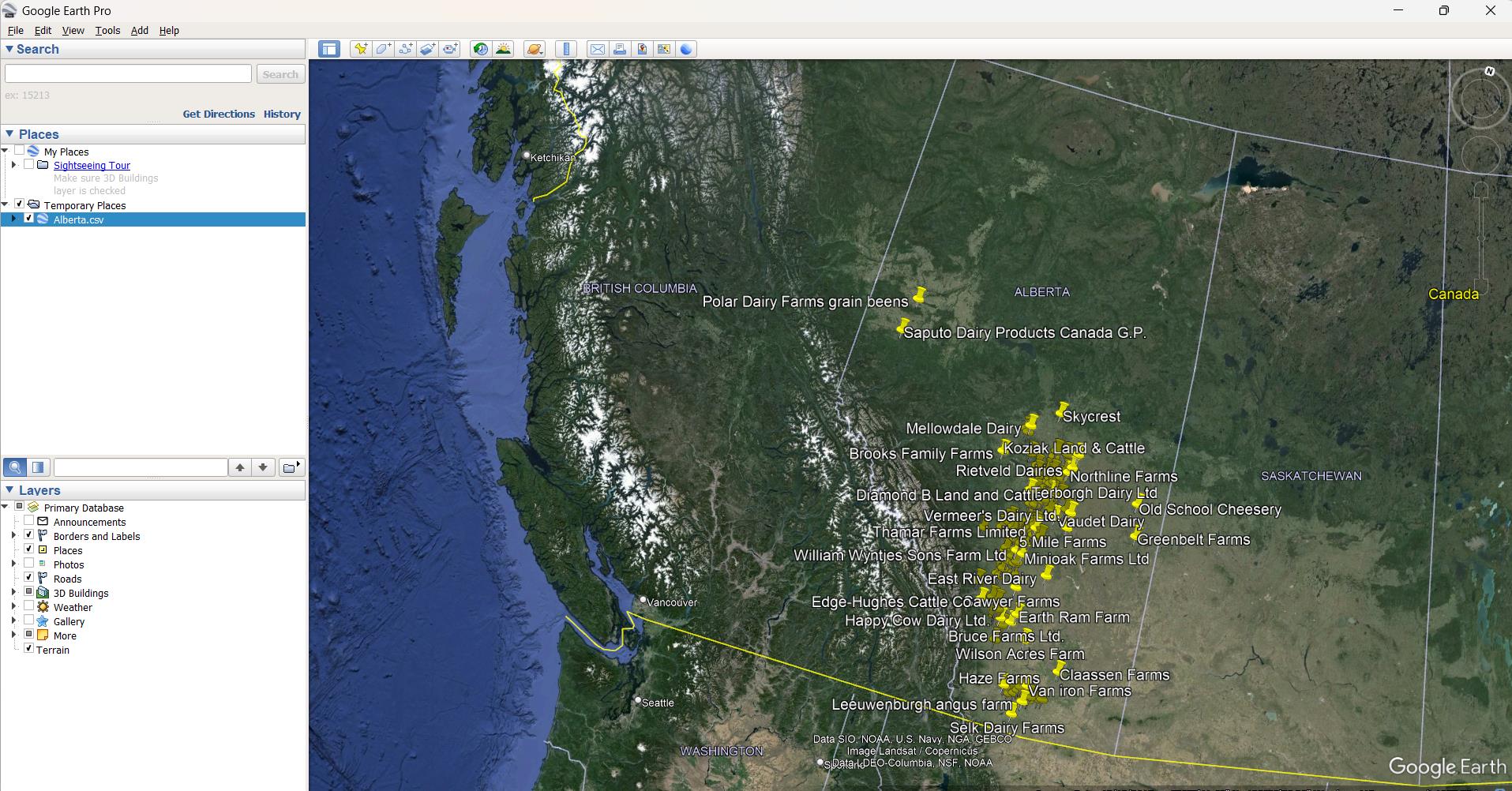

### Figure S3.png

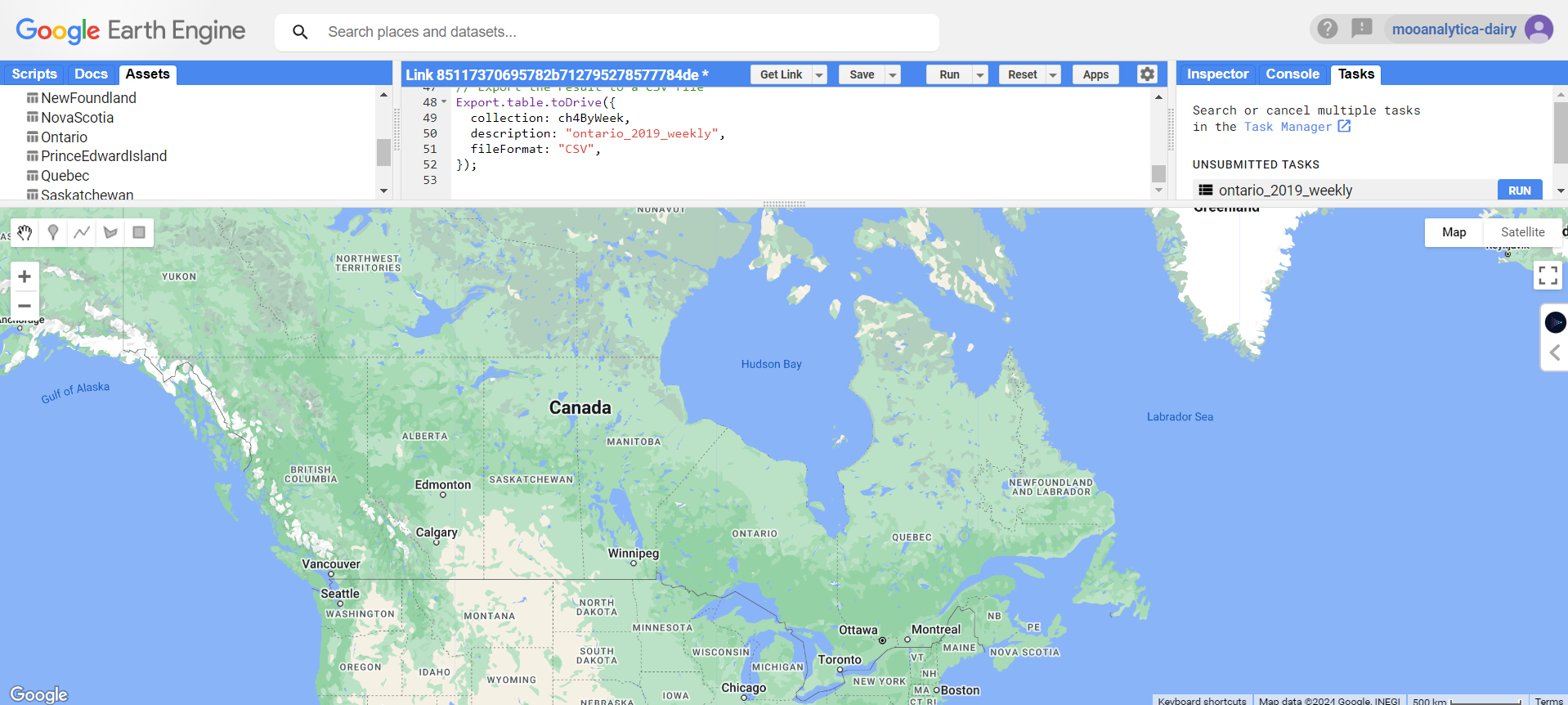
